# Genomics for reproduction in Anas platyrynchos-a novel report

**DOI:** 10.1101/2022.05.29.493861

**Authors:** Manti Debnath, Aruna Pal, Argha Chakraborty, Subhomoy Pal, Abantika Pal

## Abstract

Anas platyrynchos (ducks) are reared mostly for egg, which are very nutritious, that fetch better prices, however duck meat possess rich nutrient content. They possess the unique characteristics of disease resistance to the common avian diseases, even asymptomatic to avian influenza, with a scopeto evolve as one of the best poultry species The major limitation encountered is the lowered average egg production as well as higher age at first egg (an indicator for sexual maturity) for the indigenous ducks compared to that of exotic ones. In this current study, we attempt to explore the genes responsible for duck reproduction in terms of sexual maturity,egg production and fertility of the ducks. We had compared the genomic constitution for the Bengal duck with highest egg production with that of non-layer (infertile or sexually immature) ducks. We characterized the genes in indigenous ducks from ovarian tissues, identified important domains for characterized genes for the first time, and studied differential mRNA expression profiling for these genes with respect to layer and non-layer groups. Upregulation was observed for ESR2, DIAPH2, KMT2E, ASCF2 genes for Bengal duck in highest egg producing duck in comparison to non-layer duck, whereas downregulation was observed for KSR1, A2M, BMPR1B, ACVR1. In the next step, we explored the association with the genes which were actually responsible for egg production. Thus, duck may be utilized as a model for studying the molecular aspect of reproduction. Genes upregulated may be utilized for *knock in* of gene, whereas down regulated genes may be *knocked out* or *knocked down* through gene editing technologies for the improvement of reproductive performance of the duck in future. Molecular biomarkers may be developed with these genes for early selection of better reproducing ducks at day of hatch or even earlier.

## Introduction

Ducks have been regarded as the second most important poultry species after chicken. They are being reared for mainly egg production (layer duck) and also meat production (broiler duck) in some places. One of the important and interesting fact for duck eggs are that they have some unique features that they are more nutritious with better egg size and egg weight.

It has already been reported that ducks have a unique feature of better resistance to commonly occuring avian diseases. A remarkable feature of indigenous ducks has been observed that they are even better resistant to avian influenza virus with explored molecular basis^**1**^. Immune response genes were identified in Bengal duck providing host resistance against duck plague^**2**^. Biochemical and haematological characterization has already been studied by our group^**3**^. We had equally studied disease resistance potential for indigenous chicken Haringhata black^**4**^. We had already studied the overall growth, biomorphometry for Bengal duck in different agroclimatic region^**5,6**^. Molecular characterization of mucin gene was observed to play a unique role in growth of duck^**7**^. One of the major constraints in rearing indigenous ducks is that their egg production potential is not always economically viable. Hence our greatest challenge is to improve the egg production for them. Wide variability has been observed among egg production traits among the indigenous duck population, which opens up the scope for selection of the indigenous ducks with better egg production. It is not economically feasible to select the ducks during laying period. In case of avian specieslike duck, we consider AFE (Age at first egg) to be a greatly variable trait ranges from 125-150, average being 130±4.56 for Bengal duck^**6**^. It is not possible to determine puberty and fertility separately. Hence it is essential to develop strategies for selection of better egg producing ducks as well as early maturing and ducks with better fertility at day old stage. Molecular marker is best suited for this purpose.

Some of the important and promising genes responsible for egg production, which manifest both puberty and fertility are Protein diaphanous homolog 2 (DIAPH2), Lysine methyltransferase 2E (KMT2E), Kinase Suppressor of Ras1 (KSR1), Esr2 (estrogen receptor 2), A2M (alpha-2-macroglobulin), BMPR1B (Bone morphogenetic protein receptor type 1B), ACVR1 gene (Activin A receptor type 1, ASCF2(Acyl-CoA Synthetase Family Member 2). ESR2 or oestrogen receptor 2 has been observed to be an important protein affecting reproductive structure development. ESR1 and ESR2 act as a go-between the process of egg laying in ducks, and that ESR2 also play additional vital role for the ovary for the period of egg-laying stages ^**7**^. For enhanced egg production estrogens is a significant hub of poultry breeding and management ^**8**^. Estrogens play a essential role in the development and upholding of normal sexual and reproductive functions^**9, 10**^. The biological action of estrogens is manifested through two high-affinity estrogen receptors, ESR1 and ESR2, which go to the family of the nuclear receptor super family, the transcription factors and are articulated at different levels in target cells. ^**11-14**^

In the development and function of the ovaries, Protein diaphanous homolog 2 (DIAPH2) gene may play a role and malfunction is responsible for premature ovarian failure^**15**^. It belongs to the diaphanous subfamily of the formin homology family of proteins. It is helpful for female gamete generation. Lysine methyltransferase 2E (KMT2E) gene is encoded as KMT2E and plays a role with an N-terminal PHD zinc finger and a central SET domain. On the other way excess production is sometimes harmful because it inhibits cell cycle progression, also associated with intellectual disability, autism, macrocephaly, hypotonia, functional gastrointestinal abnormalities and epilepsy in homo sapiens. It interacts with 17 beta-estradiol, resulting in decreased expression of KMT2E (https://rgd.mcw.edu/rgdweb/report/).

BMPs are responsible for endochondral bone formation and embryogenesis. They transduce their signals by forming heteromeric complexes of two different kinds of serine (threonine) kinase receptors as type I receptors (50-55 kD) and type II receptors (70-80 kD). Type II receptors can bind ligands even without type I receptors, but they need their respective type I receptors for the purpose of signaling. On the otherhand, type I receptors need their respective type II receptors for the purpose of ligand binding. The <u>bone morphogenetic proteins</u> (BMPs), along with other intraovarian growth factors, are responsible for regulation of follicle recruitment, dominant follicle selection, ovulation, and atresia^**16**^. The BMP signaling system possess a major intraovarian role in many species. It also includes human. It aids in the generation of transcription factors which influence proliferation, <u>steroidogenesis</u>, cell differentiation, and maturation prior to ovulation, and also the formation of corpora lutea after ovulation. In the anterior pituitary, BMPs also aids to the regulation of gonadotrophin production^**16**^. NOTCH and bone morphogenetic protein (BMP)/SMAD signaling play key regulatory roles in mammalian ovarian development^**17**^. The mutation Q249R (glutamine→arginine) in the regulatory serine/threonine kinase domain of BMPR1B receptor was identified in Booroola ewes and causes increased fertility^**18**^.

A2M gene is a cytokine transporter and protease inhibitor and also uses bait-and-trap mechanism to inhibit a broad spectrum of proteases, including trypsin, thrombin and collagenase. Also known for the inhibition of inflammatory cytokines i.e., inflammatory cascades. Alpha 2 macroglobulin (A2M or ovostatin) is a homotetrameric protein consists of four disulfide-linked subunits. It has the property of inactivating/inhibiting most known proteases including serine-, threonine-, cysteine-, aspartic- and metalloproteases. Identification and biochemical characterization of chicken A2M have been undertaken, but its functional role(s) in the oviduct, hormonal regulation of expression or its expression in ovarian carcinomas in chickens is still not understood properly. A2M is novel estrogen-stimulated gene expressed in LE and GE of the chicken oviduct and may be used for monitoring effects of therapies for ovarian cancer in laying hens ^**19**^. The ACVR1 gene, helps to organize the growth and development of the bones and muscles, derive from many tissues of the body including skeletal muscle and cartilage, provide directions for the assembly of activin receptor type-1 (ACVR1) protein, a member of a protein family also known as bone morphogenetic protein (BMP) type I receptors. ACVR1 equally functions for reproduction^**20**^. BMP receptors extent the cell membrane, so that one end of the protein can stuck inside the cell to obtain signals from exterior side of the cell and can convey them within to influence cell development as well as its function^**20**^.

ASCF2(Acyl-CoA Synthetase Family Member 2) enables medium-chain fatty acid-CoA ligase activity as well as hypothtically involved in fatty acid metabolic process and can be found in mitochondrial matrix. ACSF2 (Acyl-CoA Synthetase Family Member 2) gene encode Proteine. Among its related pathways are Metabolism and Fatty Acyl-CoA Biosynthesis^**21-25**^. Gene Ontology (GO) annotations related to this gene include ligase activity. An important paralog of this gene is ACSS1. Acyl-CoA synthases catalyze the initial reaction in fatty acid metabolism, by forming a thioester with CoA (PubMed:17762044).

Marc Therrien et al. in Drosophila identified and characterized two genes, this activated Ras products are vital for transmitting signal. It doesn’t play a role in activated RAF signal. One encodes the p subunit of type I geranylgeranyl transferase, a prenylation enzyme crucial to trigger RAS to the plasma membrane. The other encodes a protein kinase that they had named kinase suppressor of ras (ksr). They also showed that this KSR has multiple receptor tyrosine kinase pathways. In their study they had isolated mammalian homologs of KSR with the Drosophila gene, which act as a novel class of kinases. Theiroutput suggest that KSR has a general and evolutionarily conserved component of the RAS signaling pathway that acts between RAS and RAF ^**26**^.RAS acting a key role in diversecellular processes for proliferation and differentiation, is responsible for nodal point transmitting signals which originates from receptor tyrosine kinases (RTKs) to a array of effector molecules^**27,30**^. Mitogen-activated protein kinase pathways are drawn in the regulation of cell differentiation, even though their precise roles in loads of differentiation programs remain elusive. The Raf/MEK/extracellular signal-regulated kinase (ERK) kinase cascade, promote and inhibit adipogenesis. In their study they had titrate the expression of the molecular scaffold kinase suppressor of Ras 1 (KSR1) to standardize signaling through the Raf/MEK/ERK/p90 ribosomal S6 kinase (RSK) kinase cascade and showed how it determines adipogenic potential. Adipogenesis can be prevented In vitro by the deletion of KSR1, which can also rescue by giving low levels of KSR1. They also showed that exact levels of KSR1can coordinate ERK and RSK activation with C/EBPbeta synthesis can lead to the phosphorylation and stabilization of C/EBPbeta within the adipogenic program. Decreased levels of KSR1 expression can enhance cell proliferation. Molecular scaffold can change the intensity and interval of signaling emanating from a single pathway to dictate cell fate, this titration of KSR1output. Kinase suppressor of Ras (KSR) is a molecular scaffold that interact with the machinery of the Raf/MEK/ERK kinase cascade and confidently regulate ERK signaling and Phosphorylation of KSR1, particularly at Ser(392) which is a critical regulator of KSR1subcellular localization as well as ERK activation. The job responsibility of phosphorylation of equally Ser(392) and Thr(274) in regulating cell proliferation and ERK activation was reported. At that time they had hypothesized that KSR1 phosphorylation is involved in activiting signaling specificity by the help of Raf/MEK/ERK kinase cascade stimulated by different growth factors. In fibroblasts, platelet-derived growth factor stimulation induces sustained promotes S-phase entry and ERK activation^**26-30**^. Earlier in our lab, we have characterized certain immune response genes in Anas platyrynchos(Bengal duck)^**31-37**^.

Hence the present study is aimed at characterization of the gene involved in reproduction of Anas platyrynchos, important domain identification in predicted 3D structure, differential mRNA expression profiling for these genes w.r.t. non layer (infertile or sexually immature) and high egg laying ducks.

## Materials and method

We studied the Bengal duck maintained in experimental farm of West Bengal University of Animal and Fishery Sciences. We maintained the production records for the birds tagged with wing band.

All the experiments were conducted in accordance with relevant guidelines and regulations of Institutional Animal Ethics committee and all experimental protocols were approved by the Institutional Biosafety Committee, West Bengal University of Animal and Fishery Sciences, Kolkata.

After completion of a part of trial, the non layer or infertile ducks and very few highest producing ducks, with certain physical deformity were culled and sold. The samples of ovary (n=6) were collected from each group of non-layer (sexually immature or infertile duck, Group I) and highest laying duck (Group II) were collected. The total RNA was isolated from the ovary of Duck by Trizol method and was further used for cDNA synthesis^**38-42**^.

### Materials

Taq DNA polymerase, 10X buffer, dNTP were purchased from Invitrogen, SYBR Green qPCR Master Mix (2X) was obtained from Thermo Fisher Scientific Inc. (PA, USA). L-Glutamine (Glutamax 100x) was purchased from Invitrogen corp., (Carlsbad, CA, USA). Penicillin-G and streptomycin were obtained from Amresco (Solon, OH, USA). Filters (Millex GV. 0.22 μm) were purchased from Millipore Pvt. Ltd., (Billerica, MA, USA). All other reagents were of analytical and molecular biology grade.

### Synthesis, Confirmation of cDNA and PCR Amplification of gene

The 20□μL reaction mixture contained 5□μg of total RNA, 0.5□μg of oligo dT primer (16– 18□mer), 40□U of Ribonuclease inhibitor, 10□M of dNTP mix, 10□mM of DTT, and 5□U of MuMLV reverse transcriptase in reverse transcriptase buffer. The reaction mixture was gently mixed and incubated at 37°C for 1 hour. The reaction was stopped by heating the mixture at 70°C for 10 minutes and chilled on ice. The integrity of the cDNA was checked by PCR. To amplify the full-length open reading frame (ORF) of gene sequence, a specific primers pair was designed based on the mRNA sequences of Gallus gallus by DNASTAR software. The primers have been listed in Table 1. 25□μL reaction mixture contained 80–100□ng cDNA, 3.0□μL 10X PCR assay buffer, 0.5□μL of 10□mM dNTP, 1□U Taq DNA polymerase, 60□ng of each primer (as in Table 1), and 2□mM MgCl2. PCR-reactions were carried out in a thermocycler (PTC-200, MJ Research, USA) with cycling conditions as, initial denaturation at 94°C for 3□min, denaturation at 94°C for 30□sec, varying annealing temperature (as mentioned in Table 1) for 35□sec, and extension at 72°C for 3□min was carried out for 35 cycles followed by final extension at 72°C for 10□min.

**Table 1:**
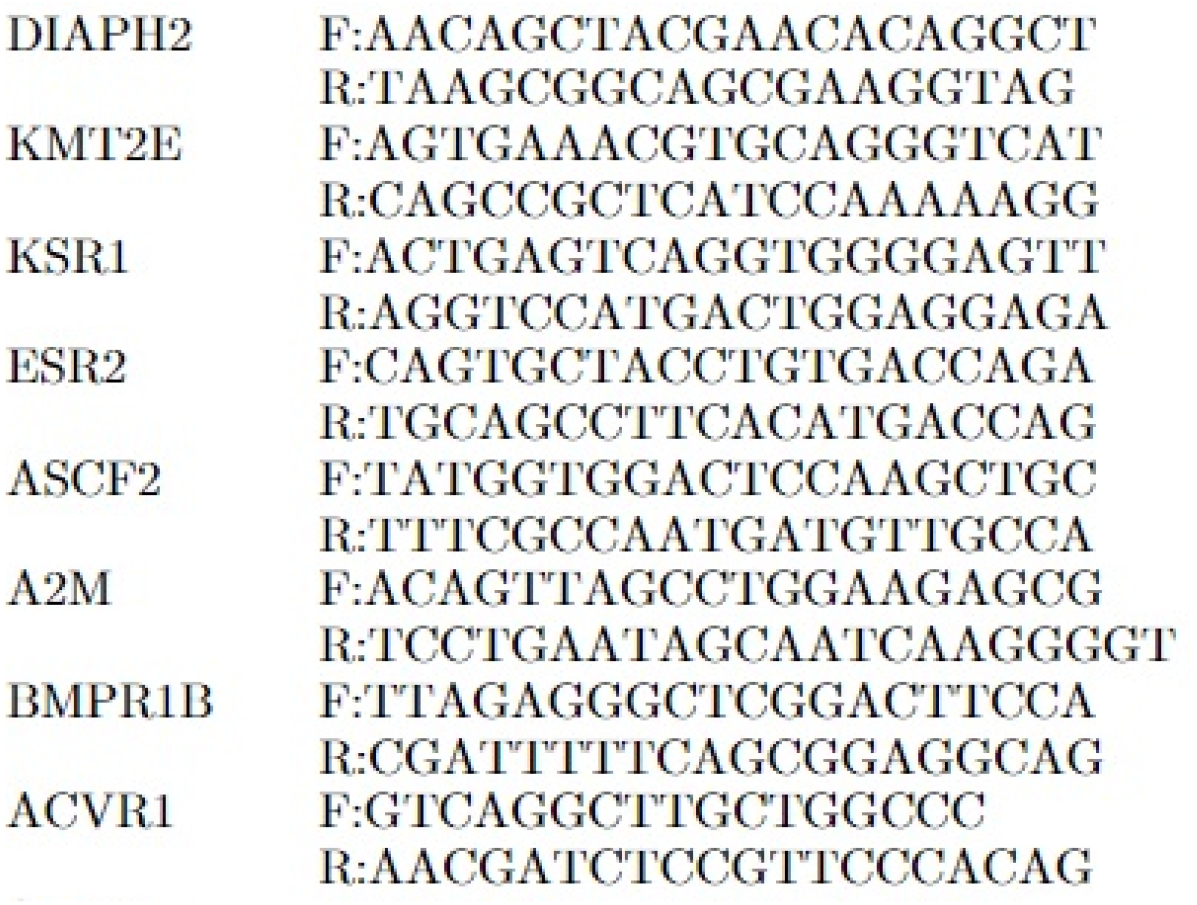
PCR primers for genes for egg traits in Duck:

### cDNA Cloning and Sequencing

PCR amplicons verified by 1% agarose gel electrophoresis were purified from gel using Gel extraction kit (Qiagen GmbH, Hilden, Germany) and ligated into a pGEM-T easy cloning vector (Promega, Madison, WI, USA) following manufacturers’ instructions. The 10□μL of the ligated product was directly added to 200□μL competent cells, and heat shock was given at 42°C for 45□sec. in a water bath, and cells were then immediately transferred on chilled ice for 5□min., and SOC was added. The bacterial culture was pelleted and plated on LB agar plate containing Ampicillin (100□mg/mL) added to agar plate @ 1:□1000, IPTG (200□mg/mL) and X-Gal (20□mg/mL) for blue-white screening. Plasmid isolation from overnight-grown culture was done by small-scale alkaline lysis method. Recombinant plasmids were characterized by PCR using gene-specific primers and restriction enzyme digestion based on reported nucleotide sequence for chicken. The enzyme EcoRI (MBI Fermentas, USA) is used for fragment release. Gene fragment insert in the recombinant plasmid was sequenced by an automated sequencer (ABI prism) using the dideoxy chain termination method with T7 and SP6 primers (Chromous Biotech, Bangalore)^**43-51**^.

### Sequence Analysis

The nucleotide sequence so obtained was analyzed for protein translation, sequence alignments, and contigs comparisons^**43-51**^ by DNASTAR Version 4.0, Inc., USA. The novel sequence was submitted to the NCBI Genbank and accession number was obtained which is available in public domain now.

### Study of Predicted gene Using Bioinformatics Tools

The predicted peptide sequence of genes of indigenous duck was derived by Edit sequence (Lasergene Software, DNASTAR) and then aligned with the peptide of other chicken breed and avian species using Megalign sequence Programme of Lasergene Software, DNASTAR (32-34). Prediction of the signal peptide of the genes were conducted using the software (Signal P 3.0 Sewer-prediction results, Technical University of Denmark). Estimation of Leucine percentage was conducted through manually from the predicted peptide sequence. Di-sulfide bonds were predicted using suitable software (http://bioinformatics.bc.edu/clotelab/DiANNA/) and by homology search with other species.

Protein sequence-level analysis study was carried out with specific software (http://www.expasy.org./tools/blast/) for determination of leucine-rich repeats (LRR), leucine zipper, N-linked glycosylation sites, detection of Leucine-rich nuclear export signals (NES), and detection of the position of GPI anchor. Detection of Leucine-rich nuclear export signals (NES) was carried out with NetNES 1.1 Server, Technical University of Denmark. Analysis of O-linked glycosylation sites was carried out using NetOGlyc 4 server (http://www.expassy.org/), whereas the N-linked glycosylation site was detected by NetNGlyc 1.0 software (http://www.expassy.org/). Detection of Leucine-zipper was conducted through Expassy software, Technical University of Denmark^**52**^. Regions for alpha-helix and beta-sheet were predicted using NetSurfP-Protein Surface Accessibility and Secondary Structure Predictions, Technical University of Denmark^**53**^. Domain linker prediction was done according to the software developed^**54**^. LPS-binding site^**55**^, as well as LPS-signaling sites^**56**^, were predicted based on homology studies with other species polypeptide.

### Three-dimensional structure prediction and Model quality assessment

The templates which possessed the highest sequence id entity with our target template were identified by using PSI-BLAST (http://blast.ncbi.nlm.nih.gov/Blast). The homology modeling was used to build a 3D structure based on homologous template structures using PHYRE2 server^**57**^. The 3D structures were visualized by PyMOL (http://www.pymol.org/) which is an open-source molecular visualization tool. Subsequently, the mutant model was generated using PyMoL tool. The Swiss PDB Viewer was employed for controlling energy minimization. The structural evaluation along with a stereochemical quality assessment of predicted model was carried out by using the SAVES (Structural Analysis and Verification Server), which is an integrated server (http://nihserver.mbi.ucla.edu/SAVES/). The ProSA (Protein Structure Analysis) webserver (https://prosa.services.came.sbg.ac.at/prosa) was used for refinement and validation of protein structure^**58**^. The ProSA was used for checking model structural quality with potential errors and the program shows a plot of its residue energies and Z-scores which determine the overall quality of the model. The solvent accessibility surface area of the genes was generated by using NetSurfP server (http://www.cbs.dtu.dk/services/NetSurfP/) ^**59**^. It calculates relative surface accessibility, Z-fit score, the probability for Alpha-Helix, probability for beta-strand and coil score, etc. TM align software was used for the alignment of 3 D structure of IR protein for different species and RMSD estimation to assess the structural differentiation^**60**^. The I-mutant analysis was conducted for mutations detected to assess the thermodynamic stability. Provean analysis was conducted to assess the deleterious nature of the mutant amino acid. PDB structure for 3D structural prediction of gene for duck was carried out through PHYRE software^38^. Protein-protein interaction have been studied through String analysis^**61**^.

### Real time PCR

Total RNA was estimated from ovary of duck from layer and non-layer group by Trizol method and quantitative analysis of total RNA were performed using formaldehyde gel electrophoresis. The 28S rRNA and 18S rRNA demarcated the quality of RNA. First strand cDNA was synthesised by the process of reverse transcriptase polymerase chain reaction (rt-PCR) in the automated temperature maintained thermocycler mechine. M-MLVRT (200 u/μl) was used as reverse transcriptase enzyme. All the primers were designed using primer 3 software (v. 0. 4.0) as per the recommended criteria. The primers used are listed in Table1. Equal amount of RNA (quantified by Qubit fluorometer, Invitrogen), wherever applicable, were used for cDNA preparation (Superscript III cDNA synthesis kit; Invitrogen). All qRT-PCR reactions were conducted on ABI 7500 fast system. Each reaction consisted of 2 μl cDNA template, 5 μl of 2X SYBR Green PCR Master Mix, 0.25 μl each of forward and reverse primers (10 pmol/μl) and nuclease free water for a final volume of 10 μl. Each sample was run in duplicate. Analysis of real-time PCR (qRT-PCR) was performed by delta-delta-Ct (ΔΔCt) method ^**31,2,4,38,39,42**^.

The entire reactions were performed in triplicate (as per MIQE Guidelines) and experiment repeated twice, in 20μl reaction volume, using FastStart Essential DNA Green Master (Himedia) on ABI 7500 system.

### Statistical analysis

Descriptive statistics with mean and standard error were estimated through SYSTAT package for the expression level analyzed through real time PCR and presented accordingly in graph. Expression level with real time PCR was estimated as 2^-ΔΔCt^

## Result

### Gross view of Ovary from laying duck vs. non laying duck

Ducks of both Group 1 (sexual immature/infertile) (Fig 1) and Group 2 (better egg laying)(Fig 2) were being dissected and ovaries are visualized. Multiple ovulation is evident in the ovary of laying duck (Fig 2).

**Fig 1A:**
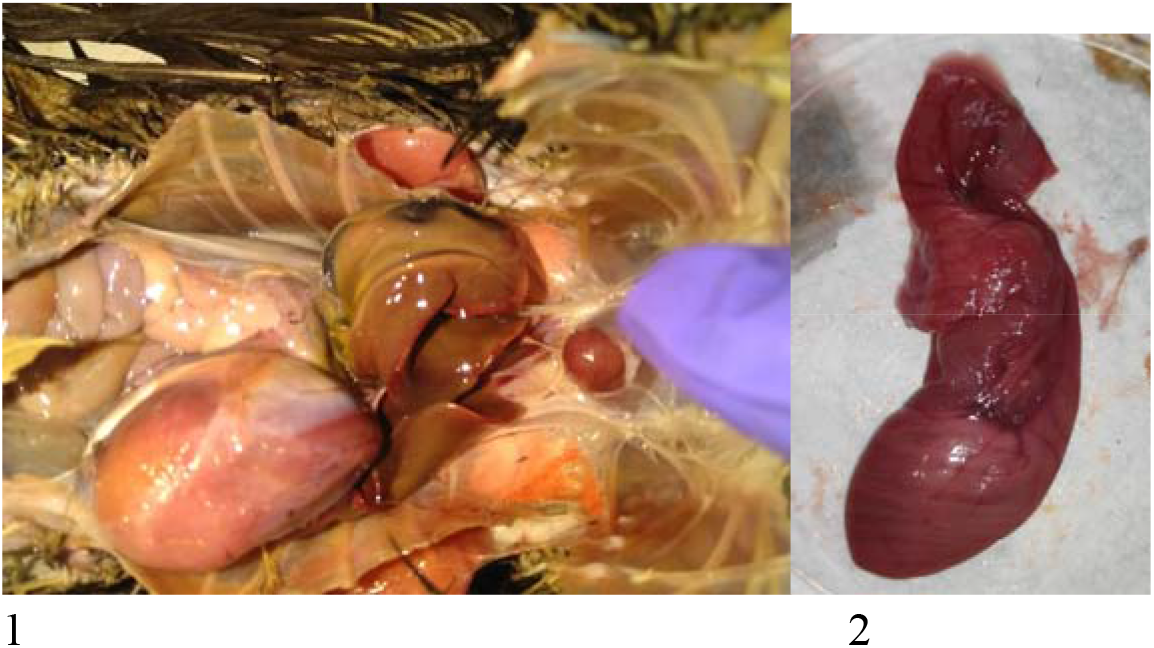
Gross Anatomical structure of ovary for *Anas platyrynchos* in non-layer

**Fig 1B:**
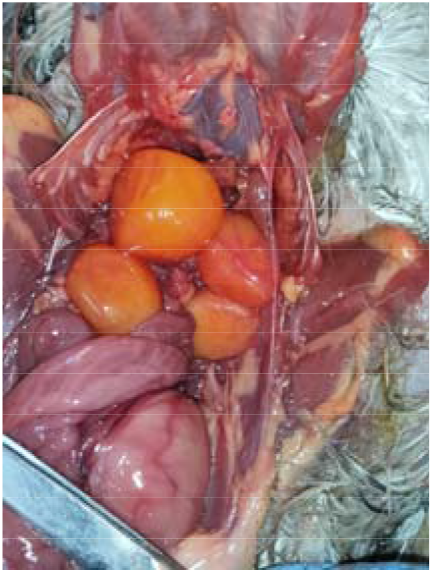
Gross Anatomical structure of ovary for *Anas platyrynchos* in layer

**Fig 2A:**
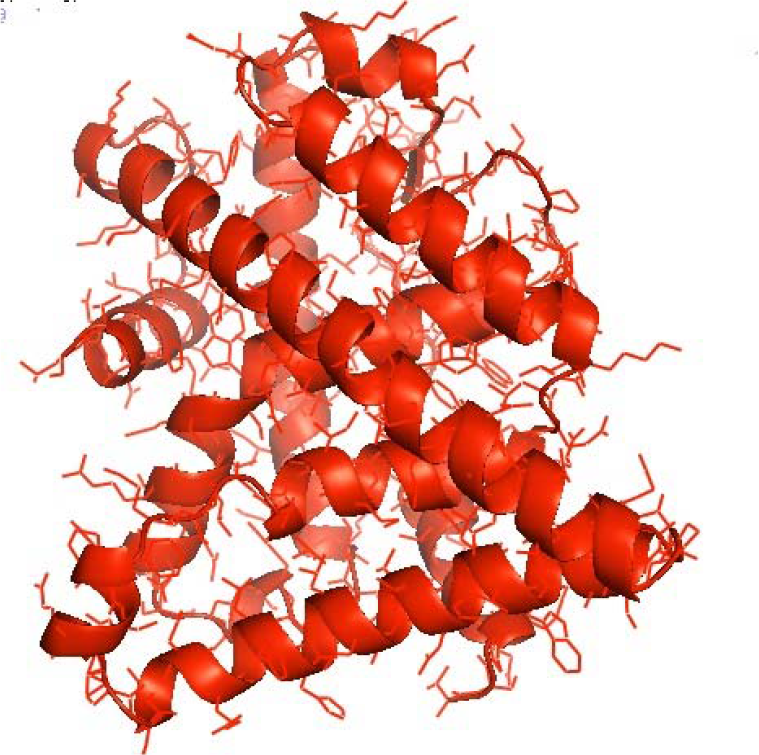
3D structure for the peptide sequence of ESR2 gene of *Anas platyrynchos*

**Fig 2B:**
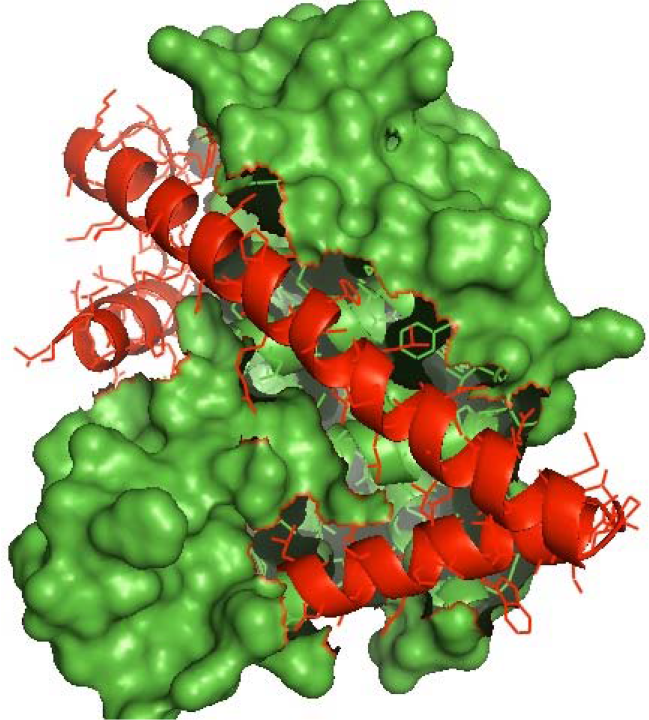
3D structure for the peptide sequence of ESR2 gene with predicted domain NR LBD at amino acid position 217-449 of Anas platyrynchos

**Fig 2C:**
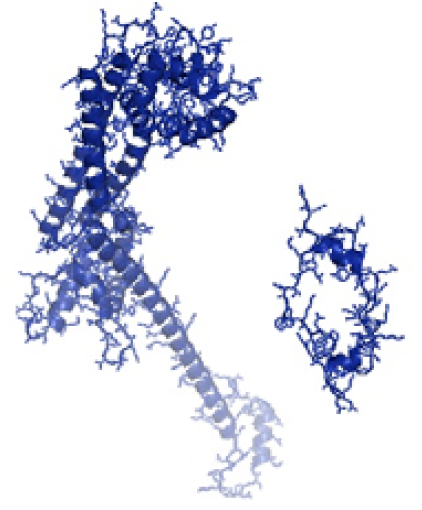
3D structure for the peptide sequence of Protein diaphanous homolog 2(DIAPH2) gene of *Anas platyrynchos*

**Fig 2D:**
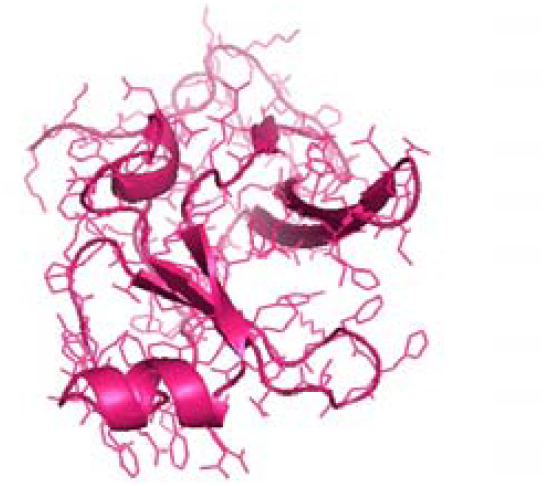
3D structure for the peptide sequence of Protein diaphanous homolog 2(KMT2E) gene of *Anas platyrynchos*

**Fig 2E:**
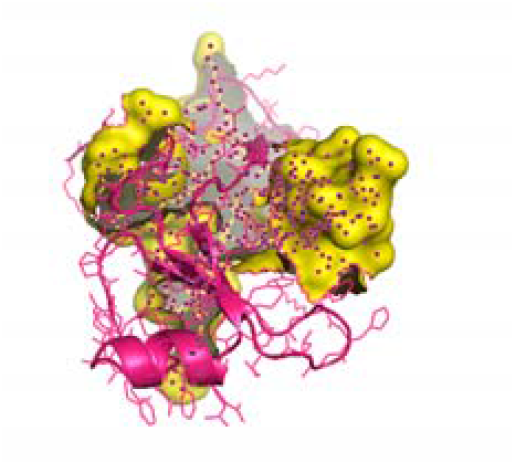
3D structure for the peptide sequence of Protein diaphanous homolog 2(KMT2E) gene of *Anas platyrynchos* with a central SET domain (330-447 a a position) yellow.

**Fig 3:**
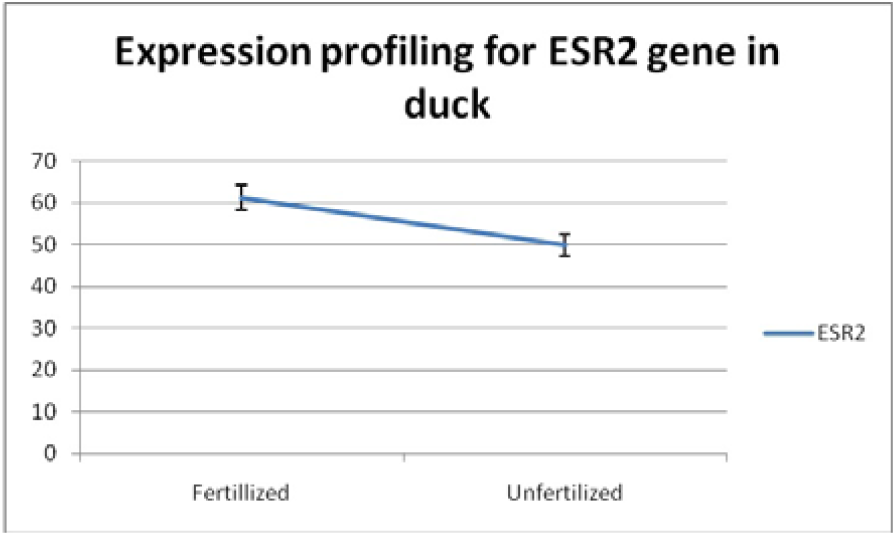
Differential mRNA expression profiling in ESR2 gene for *Anas platyrynchos* w.r.t. layer (fertilized or sexually mature) vs. non-layer (unfertilized or sexually immature)

**Fig 4A:**
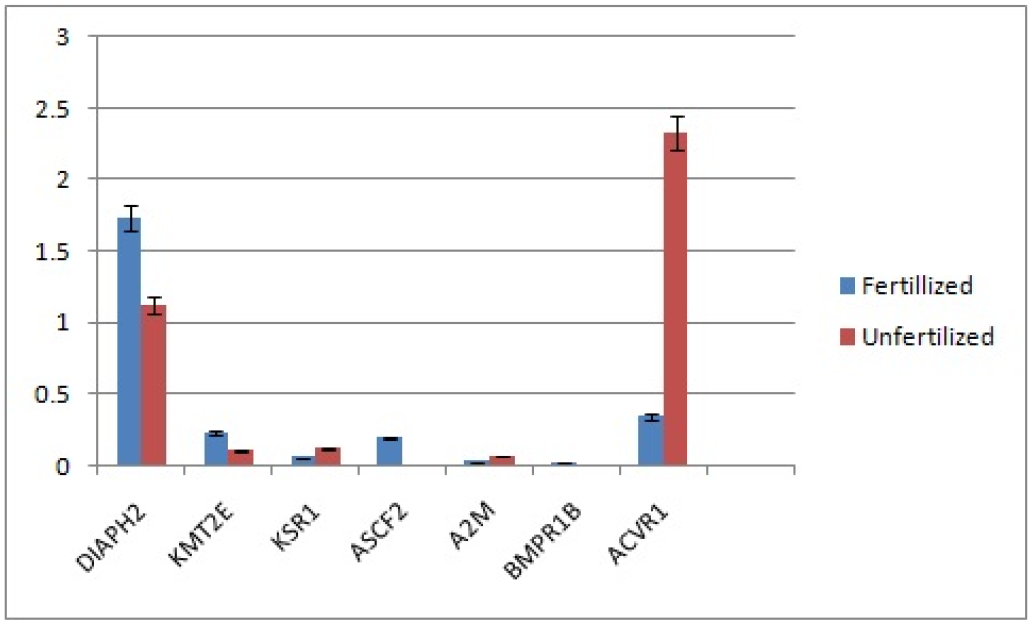
Differential mRNA expression profiling for DIAPH2, KMT2E, KSR1, ASCF2, A2M, BMPR1B, ACVR1 in *Anas platyrynchos* w.r.t. layer (fertilized or sexually mature) vs. non-layer (unfertilized or sexually immature)

**Fig 4B:**
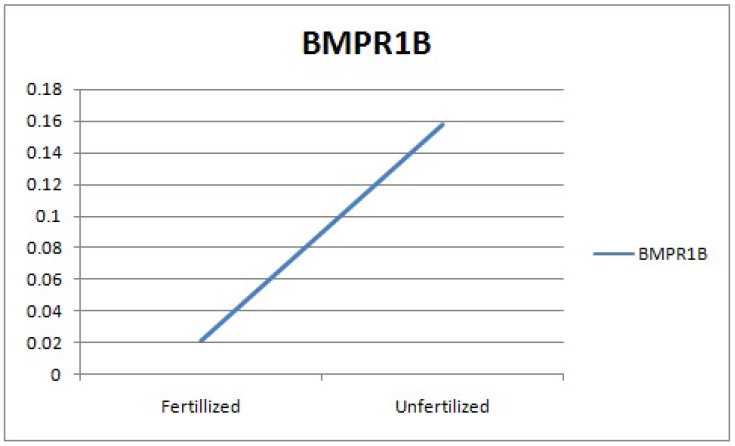
Differential mRNA expression profiling for BMPR1B in *Anas platyrynchos* w.r.t. layer (fertilized or sexually mature) vs. non-layer (unfertilized or sexually immature)

### Characterization of the genes for Sexual maturity in duck: *In silico* studies and identification of important domains

ESR2 gene has been characterized and predicted 3D structure for the peptide sequence is visualized (Fig 2A, red) and also with predicted domain NR LBD at amino acid position 217-449 (Fig 2B, green).Other important domains identified are DNA binding site as nuclear receptor had been detected at amino acid positions 105-170. Zinc finger had been detected at amino acid positions 105-125, 141-165. Modulating site for ESR2 has been identified at amino acid positions 1-104. Protein diaphanous homolog 2(DIAPH2) gene has been characterized and 3D structure predicted (Fig 2C). Lysine methyltransferase 2E (KMT2E) gene has been characterized and 3D structure predicted (Fig 2D). Important domains identified are N-terminal PHD zinc finger (amino acid position 118-166) and a central SET domain yellow coloured (330-447 a a position) Fig 2E.

### String Analysis

The analysis reports that ESR2 interacts with many other genes in the complex biological process of reproduction. These genes are NCOA1(nuclear receptor coactivator 1), NCOA3, NCOA2, SP1, ESR1, FOS, MED1,MAPK1, CAV1, NCOR2 as per their interaction score (Fig 5A, Supplementary Fig 1). The molecular interaction for DIAPH2, KMT2E, ASCF2 genes have been depicted in Fig 5B, Fig 5C and Fig 5D respectively. The detail names or abbreviation for these genes have been described in Supplementary Fig 2,3,4 with respect to these genes respectively.

**Fig 5A:**
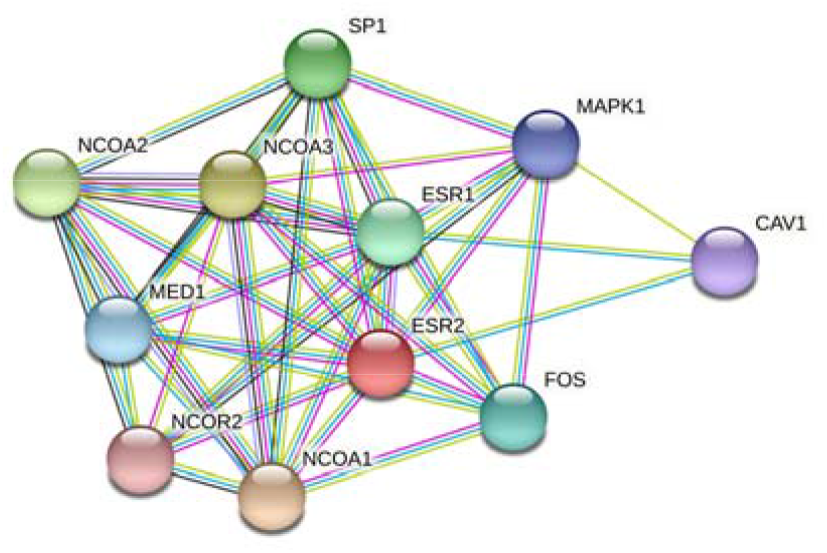
String Network analysis for ESR2 gene in *Anas platyrynchos*

**Fig 5B:**
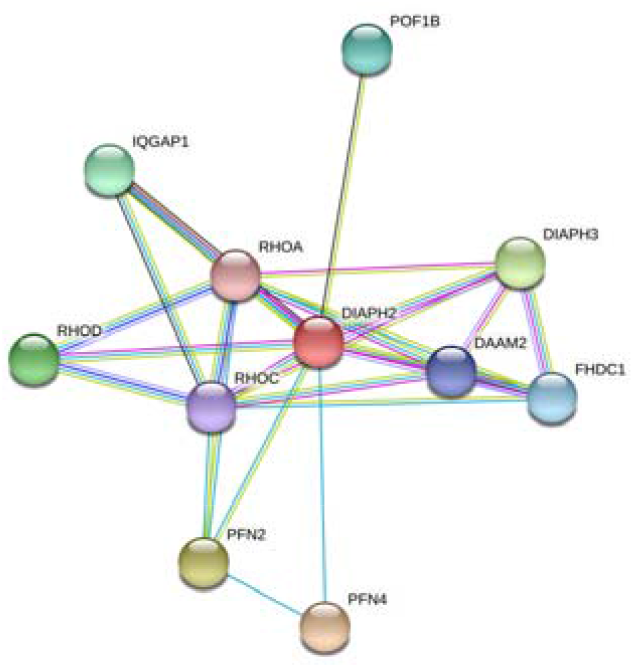
String Network analysis for DIAPH2 gene in *Anas platyrynchos*

**Fig 5C:**
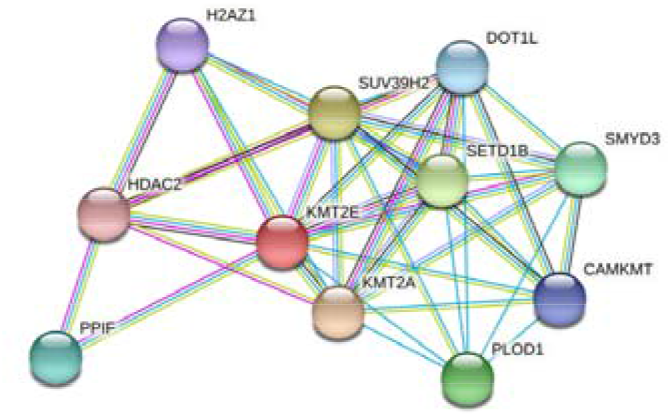
String Network analysis for KMT2E gene in *Anas platyrynchos*

**Fig 5D:**
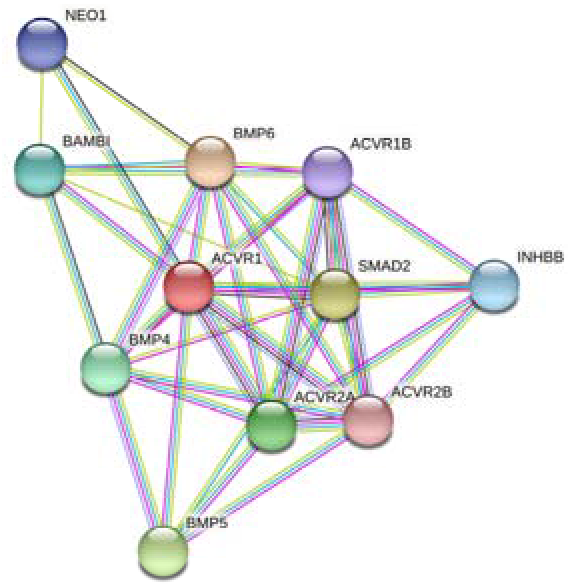
String Network analysis for ACVR1 gene in *Anas platyrynchos*

### KEGG Analysis

Role of ESR2 in estrogen signalling pathway has been described in Fig 6. It depicts the molecular pathway analysis for estrogen receptor in ovarian steroidogenesis, membrane initiated steroid signalling, nuclear initiated steroid signalling, PI3K-Akt Pathway, calcium signaling pathway.

**Fig 6:**
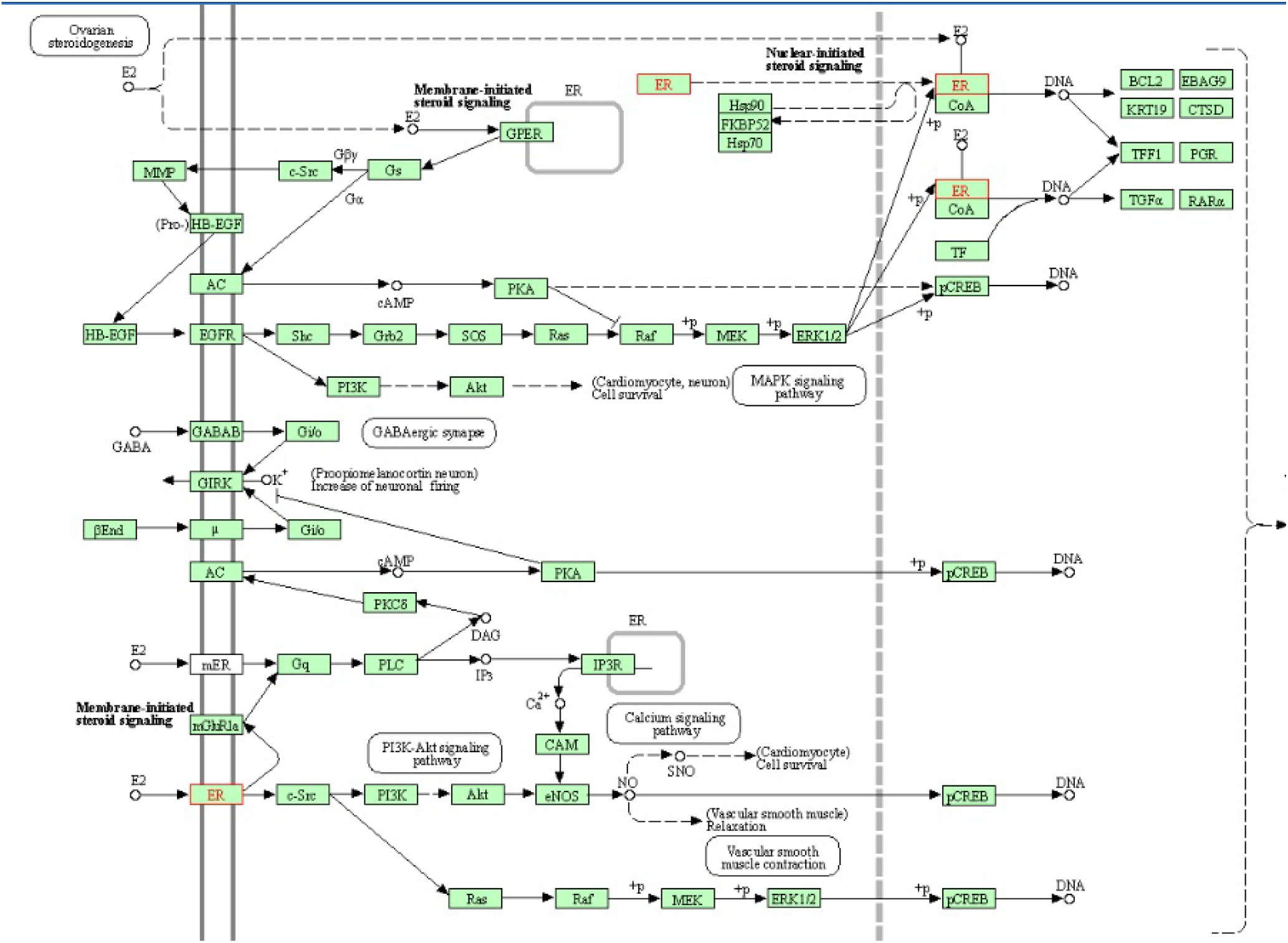
KEGG Analysis for ESR2 gene in *Anas platyrynchos*

### Differential mRNA expression profiling of the genes for egg production

ESR2 gene was observed to be upregulated in layer duck compared to that of non-layer duck (Fig 3).Upregulation was observed for DIAPH2, KMT2E, ASCF2 genes for Bengal duck in highest egg producing duck in comparison to infertile duck, whereas downregulation was observed for KSR1, A2M, BMPR1B, ACVR1 in good layer duck (Fig 4).

## Discussion

Ducks are considered among second most popular poultry species. We have already identified certain unique features for duck in terms of disease resistance to common avian diseases, including avian influenza. We had already explored some genes providing resistance to ducks (Pal et al). But the major constraint identified in indigenous duck is the delay in sexual maturity, and subsequent lesser egg production. It is a great challenge for duck industry. Thus there is an urgent need to identify the factors(proteins) responsible and explore them at molecular level. In the current experiment, we had studied duck population in two groups-Group1 (non-layers which may be sexually immature/ infertile/ anoestrous) and Group2 (good egg laying duck).

Certain promising genes were studied with respect to egg production and reproduction. Upregulation was observed for ESR2, DIAPH2, KMT2E, ASCF2 genes for Bengal duck in highest egg producing duck in comparison to non-layer duck, whereas downregulation was observed for KSR1, A2M, BMPR1B, ACVR1. This is the first report to study the molecular aspect of these genes in duck with respect to sexual maturity or anoestrous or infertility, which were phenotypically manifested as non-layer. Report is available regarding differential mRNA expression for these genes with respect to egg production trait in layer ducks^**62**^, but this is the first report for molecular control of reproduction in duck. They had also reported higher expression profile for ESR2, DIAPH, KMT2E for higher egg producing duck compared to that of lower producing duck^**62**^.

Among the genes studied, estrogen receptor 2 (ESR2) was observed to be one of the important molecule and gene expression was observed to be highest in comparison to other genes under study. Certain important domains were identified for ESR2 in duck. DNA binding site as nuclear receptor had been detected at amino acid positions 105-170. Zinc finger had been detected at amino acid positions 105-125, 141-165. ESR2 receptor binds with binds estrogens like ER-alpha, and activates expression of reporter genes containing estrogen response elements (ERE) in an estrogen-dependent manner (https://www.uniprot.org/uniprot). Thus the function for ESR2 may be summarised as estrogen receptor activity, nuclear receptor activity, sequence-specific DNA binding, steroid binding, zinc ion binding^**7-14**^. It has active role in cellular response to estradiol stimulus, intracellular estrogen receptor signaling pathway^**7-14**^. Another important domain identified is NR LBD at amino acid position 217-449. Modulating site for ESR2 has been identified at amino acid positions 1-104^**7-14**^. Thus duck ESR2 is Composed of three domains: a modulating N-terminal domain, a DNA-binding domain and a C-terminal ligand-binding domain.

Another important molecule is Protein diaphanous homolog 2(DIAPH2), whose expression was observed to be better in group II in comparison to Group I, reflecting a greater influence in promoting reproducive ability in duck. Protein diaphanous homolog 2(DIAPH2) gene may play a role and malfunction is responsible for premature ovarian failure^**15**^. It belongs to the diaphanous subfamily of the formin homology family of proteins. It is helpful for female gamete generation. It is helpful for binding to monomeric or multimeric forms of actin, including actin filaments. It also aids in actin filament organization. and small GTPase binding. It is helpful for Generation of the female gamete; specialised haploid cells produced by meiosis and along with a male gamete, thus taking part in sexual reproduction. An important domain identified is Drf_GBD from amino acid 66-148. Actin cytoskeleton through reorganization is responsible for functioning of the cell cortex, including motility, adhesion, and cytokinesis.

The assembly of the cytoskeletal components at cortical sites is regulated dynamically in a temporal and spatial manner. It has been reported that the formin family proteins aids in the reorganization of the cytoskeleton. A bifunctional autoinhibitory domain (GBD) interacts with and is regulated by activated Rho family members. Mammalian Drf3 possess a CRIB-like motif within its GBD to bind to Cdc42. Cdc42 is an essential factor to activate and guide Drf3 towards the cell cortex in order to remodel the actin skeleton^**15**^. Since DIAPH2 is directly involved with ovarian function, in our case of duck ovary, improper reproductive performance was observed for less expression of DIAPH2 gene.

We observed down regulation for this gene. ACVR1 is significantly expressed more in poor reproducing individuals. Similar result was also observed for ducks producing less number of eggs^**62**^. ACVR1 participates in several growth and reproduction related GO BPs and has well documented as involvement in biological phenomena. ACVR1 encodes for a bone morphogenetic protein (BMP) type I receptor of the transforming growth factor-beta (TGF-β) superfamily which plays a key role in cell growth while regulates several reproductive processes (such as follicular development and ovulation)^**63**^. In addition, ACVR1 regulates reproduction via the BMP and anti-Müllerian hormone (AMH) signaling^**64**^. In chickens, AMH is required for the urogenital development and germ cell migration^**65**^, is presented in early development of follicles and is expressed in small follicles^**66**^. So far, the chicken ACVR1 gene has been suggested as a positional candidate gene for body weight^**67**^, has a regulatory role during skeletal development in osteogenesis and chondrogenesis^**68**^and is expressed within the chicken granulosa and thecal layers during ovarian follicle development^**69**^.

Very few studies have been reported for genomic control of ducks for egg production or reproduction in other genes^**62,70**^. Certain other reports for molecular study for egg production or reproduction in chicken are also evolving^**71**^. Reproduction being a polygenic trait, it is being regulated by multiple interacting genes through many interacting molecular pathways. Molecular interaction of ESR2 gene had been studied and observed to interact with NCOA1(nuclear receptor coactivator 1), NCOA3(nuclear receptor coactivator 3), NCOA2(nuclear receptor coactivator 2), SP1(SP1 transcription factor), ESR1(estrogen receptor 1), FOS (Fos proto-oncogene, AP-1 transcription factor subunit), MED1(mediator of RNApolII transcripion subunit 1),MAPK1(Mitogen activated protein kinase), CAV1(caveolin), NCOR2(nuclear receptor co repressor 2) as per their interaction score in descending order. In a similar way, molecular interaction of DIAPH2 has been studied with PFN4, PFN2, DIAPH3, RHOD, IQGAP1, POF1B, FHDC1,DAAM2, RHOC, RHOA. Molecular interaction has also been studied for KMT2E, ACVR1 genes. They interact with many other proteins in the biochemical pathways involved in their functional activities. ACVR1 gene was observed to interact with bone morphogenetic protein (BMP). Pleiotropy is a common phenomenon for such genes, particularly BMP genes^**72,73**^.

## Conclusion

Delayed sexual maturity or anoestrous in indigenous duck is a major problem in rearing Bengal duck, although they exhibit greater disease resistance potential. We could explore ESR2, DIAPH2, KMT2E as the prominently explored gene to be upregulated in good layer duck, whereas ACVR1 to be down regulated. Specific gene edited good laying ducks may be developed with knocked in gene (ESR2, DIAPH2, KMT2E), or knocked out or knocked down gene for ACVR1.

## Acknowledgement

The authors are thankful to Department of Biotechnology, Ministry of Science and Technology, Govt. of India (Grant number BT/PR24310/NER/95/649/2017) and Department of Science and Technology, Govt. of India (Grant no. EMR/2016/003554) for providing the financial support. The technical and financial support by Vice-Chancellor, West Bengal University of Animal and Fishery Sciences is duly acknowledged. Thanks to Director, AH & VS, Animal Resource Development Department, Govt. of West Bengal.

## Figures

**Supplementary Fig 1:**
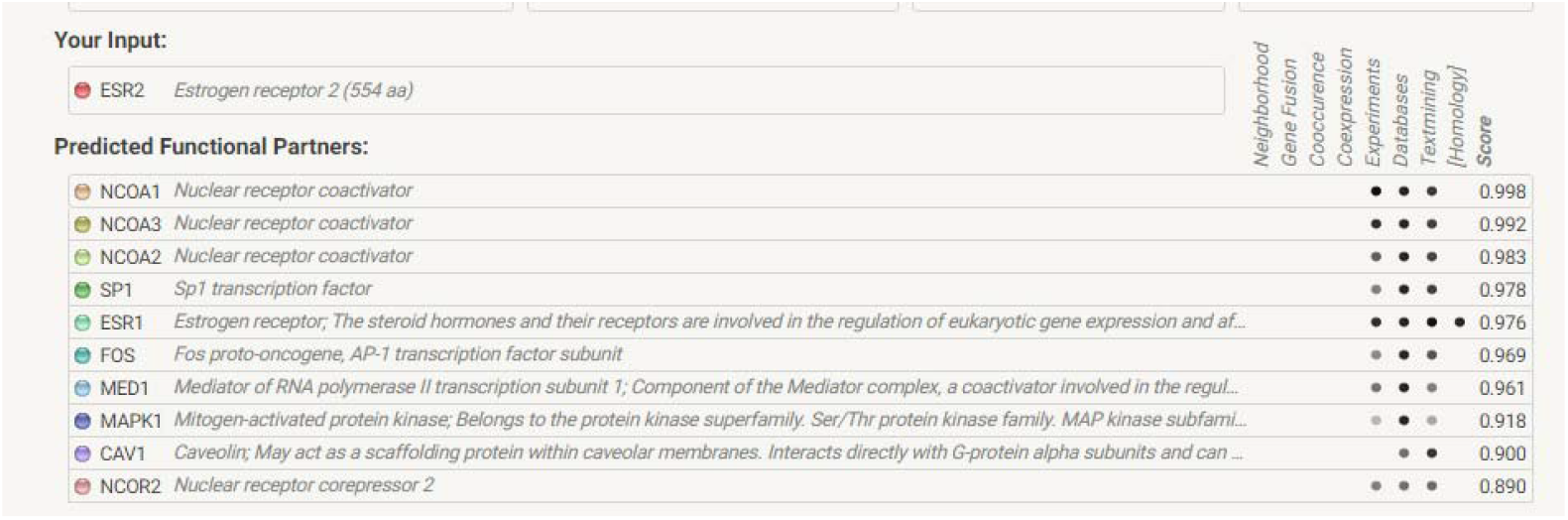
Interacting genes for *Anas platyrynchos* with ESR2

**Supplementary Fig 2:**
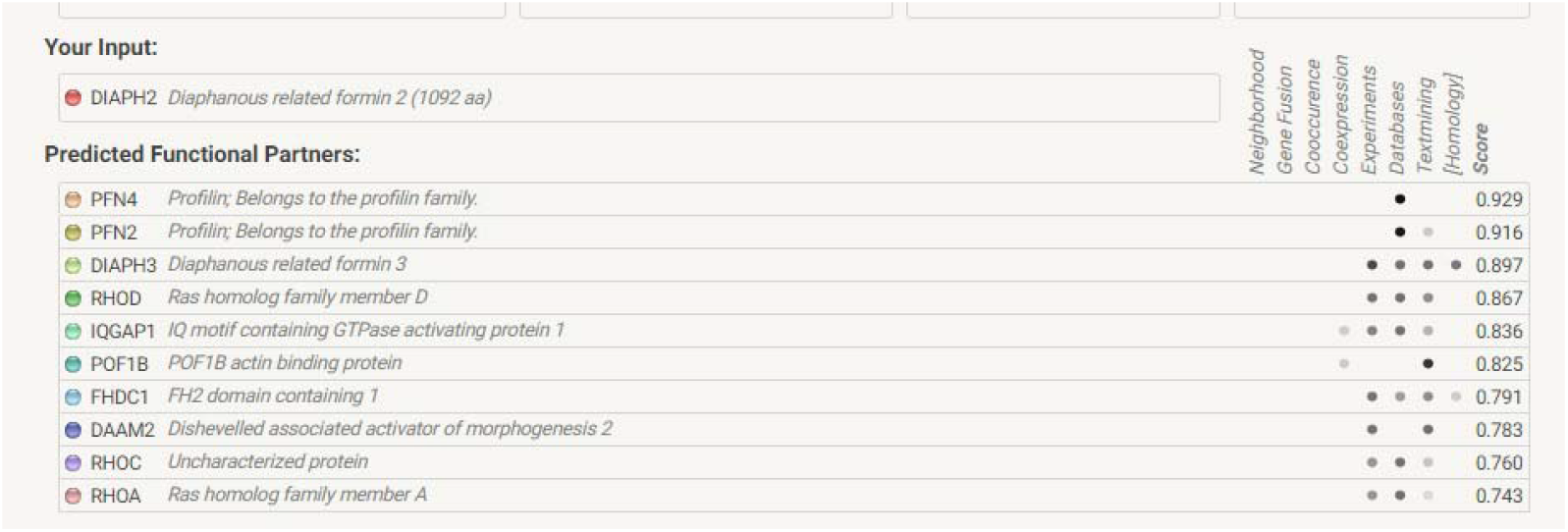
Interacting genes for *Anas platyrynchos* with DIAPH2

**Supplementary Fig 3:**
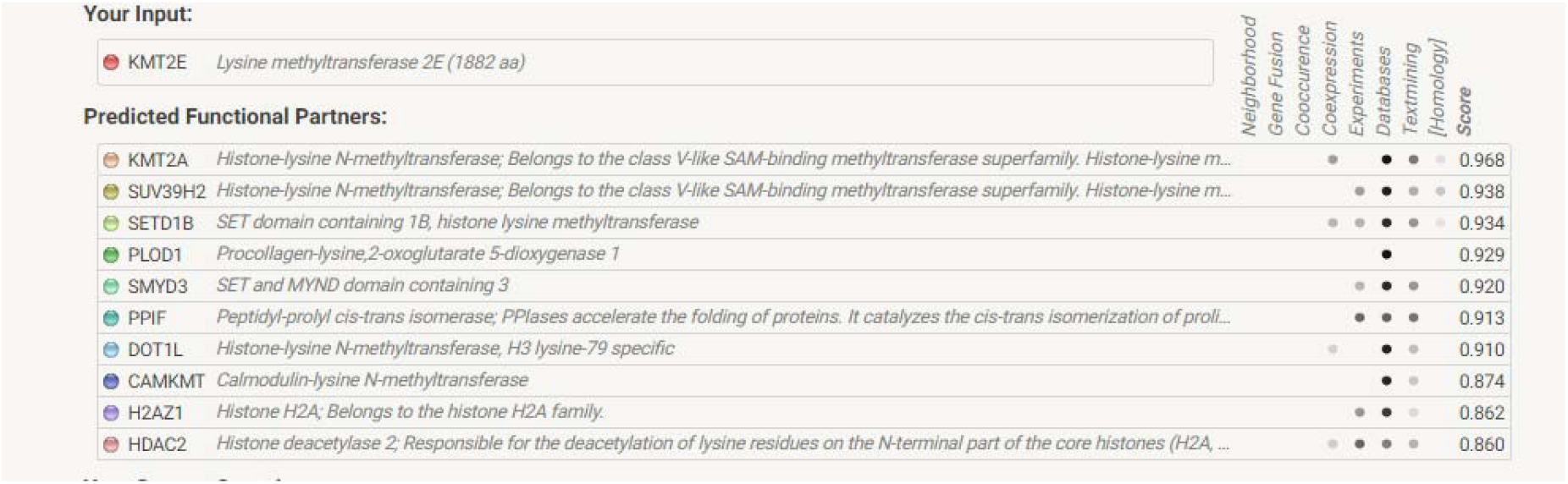
Interacting genes for *Anas platyrynchos* with KMT2E

**Supplementary Fig 4:**
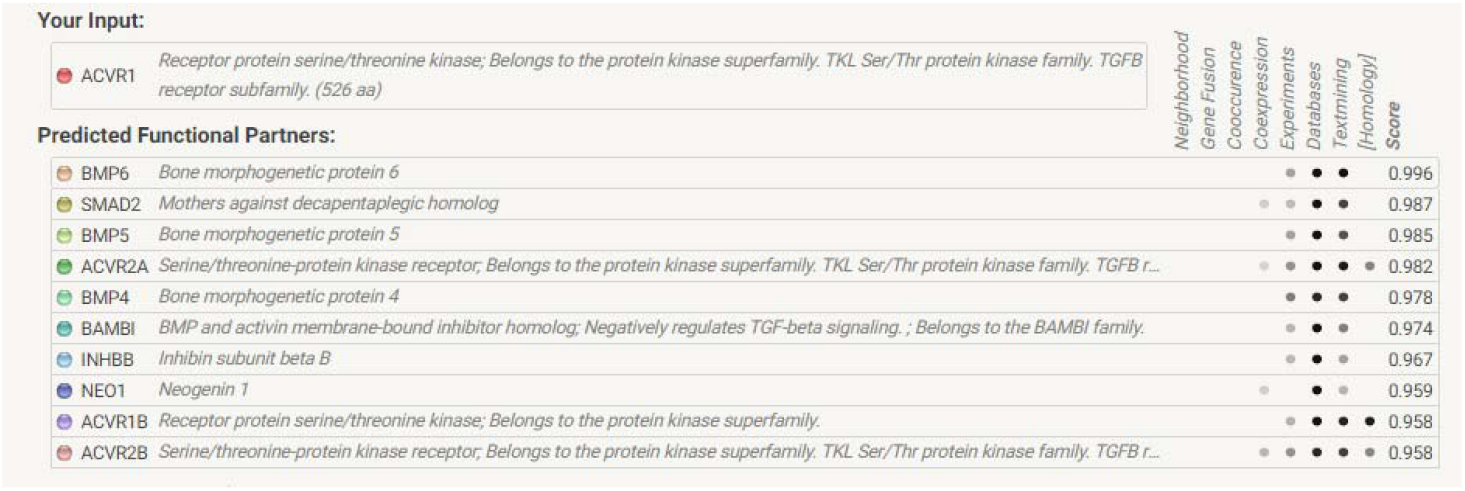
Interacting genes for *Anas platyrynchos* with ACVR1

